# Affinity-based selection of anti-Feline Leukaemia Virus p27 monoclonal antibodies for efficient lateral flow assay development

**DOI:** 10.1101/2025.09.29.679151

**Authors:** Hadyn Duncan, Elisabeth Domingo-Contreras, Sotirios Athanasiou, Rosario Fernández Godino, Francisco Castillo, Ana Camacho

## Abstract

Classic diagnostic modalities—including RT-PCR and proviral DNA PCR—present certain drawbacks, highlighting the need of alternative diagnostic techniques, such as lateral flow immunoassays (LFIA), valued for their rapidity and sensitivity. Said desirable features depend on the efficiency of the ternary interaction wherein the antigen bridges immobilized capture and labelled detection antibodies. However, the cost and the throughput of current screening methods, for finding monoclonal antibody (mAbs) pairs efficient at establishing this type of ternary interactions, remains as a universal bottleneck for LFIA development. Traditional sandwich enzyme-linked immunosorbent assays (ELISA), although extensively employed in mAb screening procedures, is burdened by its high consumption of labelled mAbs and the potential alteration of antigen conformation upon plate immobilization, often yielding suboptimal surrogates for native interactions within the final LFIA setup. In this study, we developed an orthogonal approach integrating ELISA and Spectral Shift (SST) that jointly enable the minimisation of reagent consumption (mAbs, antigens, labelling reagents) to comprehensively profile a panel of mAbs against a bona fide antigen of Feline Leukaemia Virus FeLV p27. This facilitated the quick selection of two low nanomolar mAbs with distinct epitope recognition, for which we validated a ternary complex formation in a LFIA prototype. This strategy enables advanced, resource-efficient LFIA development for FeLV and other critical antigens, such as H5 Avian Influenza Virus or African Swine Fever Virus, in addition to major human pathogens including *Helicobacter pylori* or Mpox Virus, facilitating sensitive diagnostics and streamlined quality control.

## 1. Introduction

In conventional antibody pairing approaches, sandwich ELISAs are commonly used to define binding interactions across panels of antigen-antibody trinary combinations for LFIA (lateral flow immunoassays) development (1). However, these assays often require enzymatic or fluorescently labelled detection antibodies, which can be problematic as these protocols frequently demand large amounts of expensive reagents that are often scarce in early development (2). Additionally, surface immobilization of antigens on ELISA plates can alter their native conformation and binding affinities (3), leading to results that may not accurately reflect the epitope binding event in solution and in a flowing system, potentially resulting in undesirable false positives or false negatives when detecting the antigen of interest, representing a methodological gap to be filled according to the literature in the field (4,5).

In recent years, real-time quantitative biomolecular interaction techniques, such as Isothermal Titration Calorimetry (ITC), Surface Plasmon Resonance (SPR), and emerging Spectral Shift Technology (SST) have been proposed for the determination of dissociation constants (K_d_s) between different types of interactions, including antibody-antigen complexes (6,7). These biophysical techniques allow to precisely characterize binding constants of a wide range of biomolecular associations by measuring changes in refractive index, heat exchange, or fluorescence emission variations. Even though ITC enables a more thorough characterization of the thermodynamic mechanism of binding, SPR and SST present the exclusive possibility of minimizing reagent consumption down to the microgram scale (7–10). Of the available approaches, SST can precisely characterize binding events using environment-sensitive fluorescent dyes that are attached to the antigen by means of straightforward labelling protocols, while all components, including unlabelled antibodies forming the ternary sandwich, continue in solution, in a more near-to-native manner, and without any surface immobilization requirements (10). These characteristics enable a quick method development to precisely establish competition and ternary interaction assays (11), and, as result, to immediately classify each antibody based on epitope recognition when the structural information is scarce, accelerating the selection of antibody pairs suitable for LFIA development at early discovery stages.

To fill the methodologic gaps described above, the proof of concept presented in this work is based on the Feline Leukaemia Virus (FeLV), which is considered one of the most predominant contagious feline infections to affect domestic cats and wild felines worldwide. This retrovirus is primarily transmitted through saliva and targets the felines immune system and hematopoietic cells causing immunosuppression and anaemia, often leading to lymphoma and leukaemia (12). Based on the clinical importance of FeLV, a vast number of diagnostic techniques have been made available for the determination of infections. These essential tools for FeLV disease control include traditional methods based on viral RNA detection by RT-PCR and proviral DNA PCR analysis, which present limited performance (13,14), and, alternatively, the widely adopted in the field of veterinary diagnosis, LFIA (see Figure 1 below) offering quick time to results with high accuracy by primarily targeting the highly immunogenic p27 core protein of FeLV, or FeLV p27 in blood, mucus, or serum.

**Figure 1.**
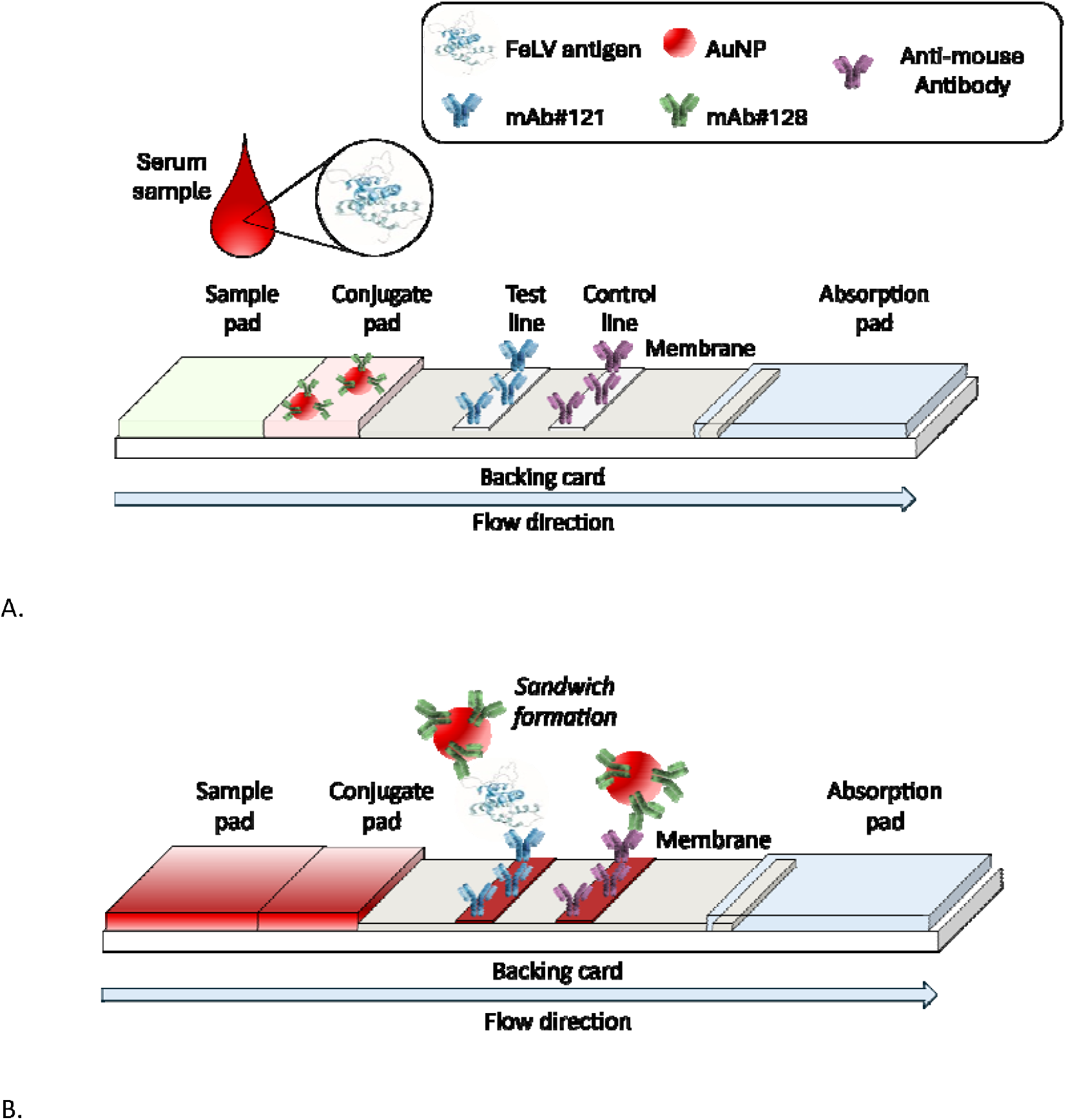
Scheme of the LFIA for FeLV p27 antigen detection via sandwich formation. (A) Serum sample containing the target antigen is applied to the sample pad, from where it is transported by capillary forces to the conjugate pad to interact with the labelled anti-FeLV p27 mAb. (B) Test flow carries the antigen-antibody complex throughout the membrane and binds to the second anti-FeLV p27 mAb immobilized in the test zone area, resulting in the appearance of a visible red line.

The FeLV p27 protein (15), which is a key part of FeLV Gag polyprotein, is a highly conserved, immunogenic capsid component that plays a central role in both viral structure and host immune recognition. p27 adopts a capsid-like, β-sheet-rich conformation with stabilizing α-helical regions, and its N-terminal domain is characterized by increased flexibility and surface exposure, theoretically facilitating immune recognition (16). FeLV p27 seemingly exhibits several key conserved immunogenic epitopes, particularly spanning residues 1–21, 86–98, 115–125, and 195–210 (16), as well as potential conformational epitopes integral to the native structure. Accordingly, a variety of mAbs have been developed to target these immunodominant regions, especially within the central and C-terminal domains (16). Owing to its conserved nature and multiple accessible epitopes, p27 serves then as a robust target for novel assay development, and even for proof-of-concept studies (17,18) to validate emerging methods and diagnostic platforms. However, despite the integration of high-quality immunoreagents in current FeLV tests, insufficient characterization of antigen-antibody interactions may result in suboptimal antibody pairing, potentially limiting the effectiveness of existing LFIA-based medical devices (2).

The present work describes a strategic selection of immunoreagents, based on orthogonal characterization by ELISA and SST of binding interactions within a panel of novel mAbs generated *ad hoc* to target FeLV p27. The method enabled the identification of two low nanomolar mAbs that bind FeLV p27 at distinct epitopes and efficiently form a ternary sandwich, with these binding characteristics subsequently tested in a LFIA prototype using serum samples containing FeLV p27. The methodology was validated at higher technology readiness levels, offering an unprecedented quick approach for sensitive and specific LFIA development for other high priority antigens of *One Health* interest (https://www.who.int/health-topics/one-health#tab=tab_1), and for quality control assays that use quantitative biomolecular recognition data while minimizing sample requirements and associated costs (19,20).

## 2. Results and Discussion

### 2.1. Monoclonal antibody production against FeLV p27 antigen

As described in the materials and methods section of this article, *in vivo* mice immune response for antibody production was generated by inoculating periodically recombinant FeLV p27 antigen. Ten-days after the third injection, antisera samples were collected and evaluated by ELISA in detection format. Results showed high sensitivity towards the FeLV p27 antigen, with half maximal effective concentration (EC_50_) values below 1 nM (Figure S1). Next, B cells isolated from immunized mice were used to generate mAbs following the hybridoma technology protocol described herein. Preliminary screening showed twenty-five parental hybridomas with positive reactivity towards the FeLV p27 antigen. After three cycles of single cell cloning procedures and ELISA analysis, four well-grown hybridoma cell lines were stabilized (#14A, #14B, #121 and #128) and specific anti-FeLV p27 mAbs were purified from culture supernatant. Finally, SDS-PAGE analysis and rapid ELISA demonstrated that the four purified mAbs were IgG_1_/κ type immunoglobulins (Figure 2). For reference, two control anti-FeLV p27 mAbs (#090 and #080, see below for further details) were included.

**Figure 2.**
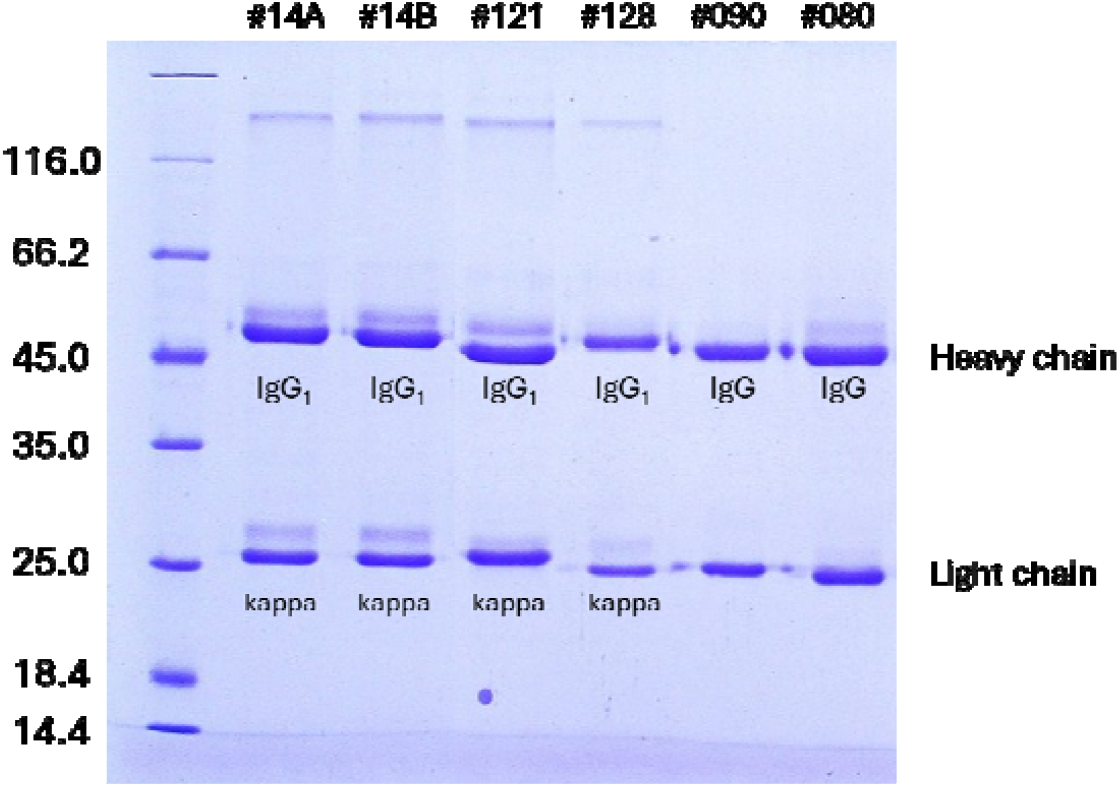
SDS-PAGE analysis of all anti-FeLV p27 monoclonal antibodies. Isotype of reference antibodies #090 and #080 is based on information made available by supplier as described in materials and methods.

### 2.2. Monoclonal antibody characterization by EL

Antibody affinity was assessed using two ELISA formats. Capture ELISA immobilised the antibody to test its ability to bind free antigen. Detection ELISA attached the antigen to the plate and checked for antibody binding in solution. Commercial mAb #090 served as the capture reference antibody. Commercial mAb #080 acted as the detection reference antibody. These choices were based on supplier information. This stepwise approach ensured a thorough and comparative analysis of antibody performance. To ensure a higher degree of epitope exposure and avoid protein denaturalization due to surface immobilization, both ELISA modalities were carried out employing a monobiotinylated form of the recombinant FeLV p27 protein, in which a biotin moiety is attached to a BCCP-tag in the C-terminus region of FeLV p27 by E. coli biotin ligase BirA (21). Biotinylation ensured a consistent orientation of antigens and improved detection sensitivity and assay reproducibility compared to other methods like direct adsorption that may cause partial epitope masking (21). As such, determination of EC_50_ as apparent affinity constant in capture ELISA were obtained from the fitting of the raw data/Absorbance values to a four-parameter logistic regression model (4PL, raw data shown in Figure 3A, and EC_50_ values summarized in Table 1). Complementarily, in detection ELISA, assay configuration was reversed allowing for affinity analysis of each antibody towards the immobilized protein (raw data shown in Figure 3B, and corresponding EC_50_ values summarized in Table 1) (22,23). Based on the EC_50_ values obtained, we performed a ranking of affinity for the mAb candidates. The new candidates #14A, #121, and #128 presented sub nanomolar to nanomolar affinities in both antigen capture and antigen detection assays, therefore a better recognition of their corresponding immunogenic epitopes, which was independent from the assay setup. The better affinity and epitope accessibility features will be expected to ensure improved sensitivity for immunoassays, including LFIAs devices (24–27).

**Table 1.** Summary table of the characterization of Monoclonal Antibody Affinities for FeLV p27. Summary of EC_50_ (nM) of six mAbs for the target antigen, as determined by two complementary techniques: Capture ELISA and Detection ELISA. Further correlation with K_d_ (nM) values obtained by Spectral Shift assay is shown. In Capture ELISA, mAbs are immobilized to assess their ability to bind soluble antigen (Ag) using a colorimetric assay. Detection ELISA involves a reversed setup which attaches the antigen on the plate surface and colorimetrically detects the mAb bound in solution. The Spectral Shift assay evaluates binary and ternary mAb–antigen interactions in solution, both with free antigen and under sandwich conditions (1 nM antigen pre-saturated with 20 nM reference mAb #090). As described in materials and methods, all EC_50_ and K_d_ values were obtained by fitting data with a four-parameter logistic regression model (4PL, Log[mAb] vs. response, variable slope) using GraphPad Prism 10 as described in materials and methods. For fits with errors exceeding 25%, EC_50_ and K_d_ values are reported as estimates.

| mAb | ELISA ($EC_{50}$, nM) | | Spectral Shift ($K_d$, nM) | | |
| --- | --- | --- | --- | --- | --- |
|  | Capture | Detection | 5 nM Ag | 1 nM Ag | 1 nM Ag + 20 nM #090, sandwich |
| #090 (ref) | ~0.22 | ~0.1 | ~2.46 | - | - |
| #080 (ref) | ~0.75 | ~0.3 | Nbd<sup>a</sup> | - | - |
| #14A | $0.08 \pm 0.02$ | $0.04 \pm 0.01$ | $0.12 \pm 0.03$ | ~0.13 | $2.01 \pm 0.05$ |
| #14B | ~0.89 | $0.06 \pm 0.01$ | $0.16 \pm 0.04$ | ~0.31 | $0.35 \pm 0.15$ |
| #121 | $0.06 \pm 0.02$ | ~0.13 | ~0.74 | ~4.64 | Nbd |
| #128 | ~0.11 | $0.07 \pm 0.02$ | $0.16 \pm 0.02$ | ~0.23 | $0.22 \pm 0.06$ |

<sup>a</sup> Nbd = No significant binding detected.

**Figure 3.**
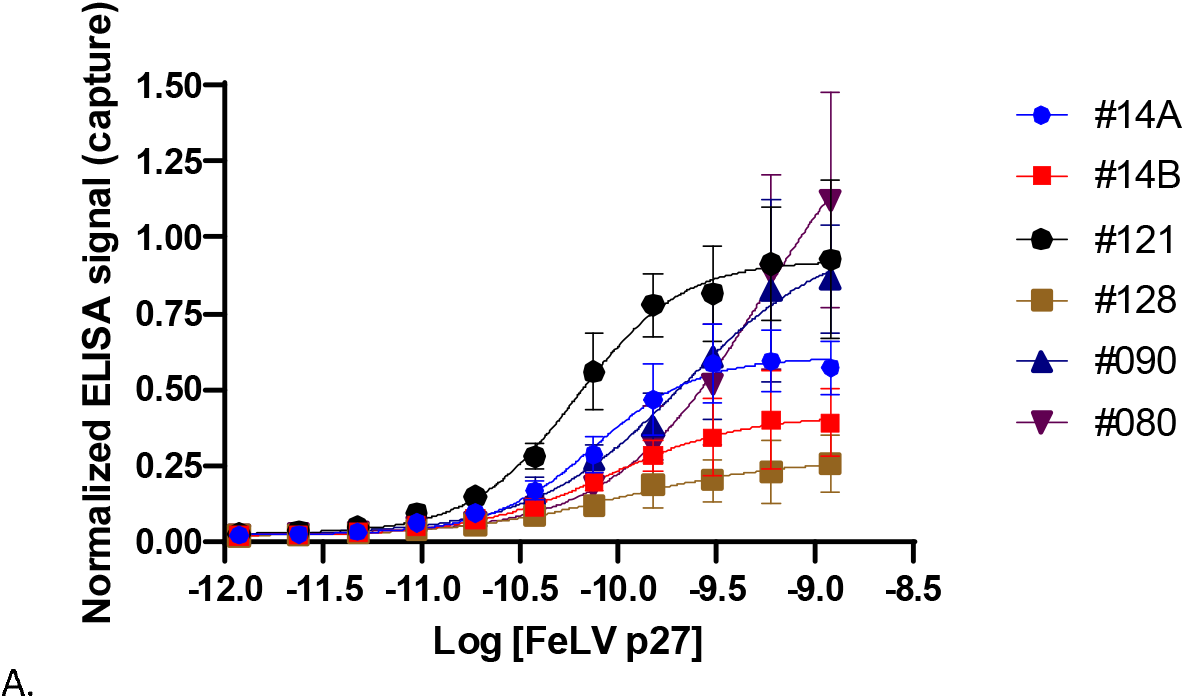
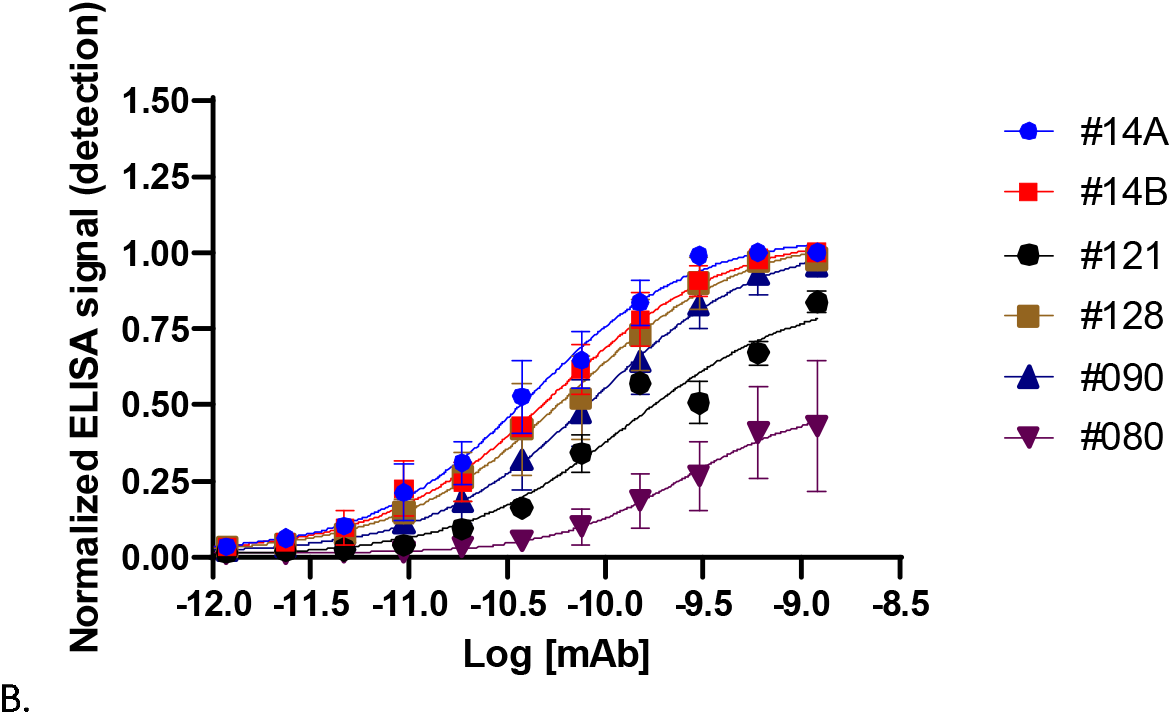
ELISA Characterization of Monoclonal Antibody Affinities for FeLV p27. A) Capture ELISA measured binding of 7.14 nM mAb (adsorbed to the plate) to serially diluted FeLV p27 antigen in solution (starting at 12 nM, x-axis in log scale). B) Detection ELISA used serially diluted mAbs in solution (from 12 nM, x-axis in log scale) with 12 nM FeLV p27 adsorbed to the plate. In both cases, Absorbance values (y-axis) reporting for mAbs/FeLV p27 binding was analysed via a 4PL model to estimate EC_50_ values summarized in Table 1.

### 2.3. Monoclonal antibody binding and antibody paring analysis by Spectral Shift Technology (SST)

Next, and to avoid any affinity or avidity artifacts coming from the ELISA plate setup (28) that could mislead antibody pairing experiments, a thorough characterization of the binding properties of selected mAbs was performed, for which a biotin-free FeLV p27 antigen labelled at its primary amines by a RED-NHS dye was used (conjugation protocol described in materials and methods section below). Accordingly, affinity and K_d_ values were measured by an SST assay, performed at two different concentrations of labelled antigen. 5 nM NHS-labelled FeLV p27 was defined as a standard conditions binding assay, to orthogonally validate ELISA results; and 1 nM NHS-labelled FeLV p27, as a stringent selection assay. The rationale of this two-stages approach was that when antibody candidates are evaluated in solution at reduced antigen concentrations, e.g. a 5-fold decrease, only those with the strongest intrinsic affinities would maintain detectable complex formation, indicative of enhanced analytical sensitivity in the final LFIA devices. Thus, affinity assays under stringent conditions could filter out low priority mAbs, while saving significant amounts of expensive reagents, and before entering more laborious experiments in the LFIA format, where the antigen is present at very low concentrations in a flowing system (28,29). The sub nanomolar to nanomolar EC_50_ values from ELISA, reporting for apparent affinity to FeLV p27, matched K_d_ values obtained in SST under standard conditions (see Figure 4A, Table 1-5 nM Ag column) for the reference #mAb 090 and the new candidate mAbs #14A-B, #121, and #128. Overall, these results validated the expected orthogonality between ELISA and SST affinity methods. Said K_d_ values from the standard SST assay also showed a consistent trend with those K_d_ values from stringent conditions for new candidates mAbs #14A-B, #121, and #128 (Figures 4B–4E, and Table 1: 1 nM Ag column), which were then defined as more sensitive/leading mAbs for further pairing experiments and epitope profiling. In this regard, reference monoclonal antibody #090 emerged as the optimal choice for setting up subsequent SST-based ternary binding assays, outperforming #080 in this role. Notably, #080 failed to show significant binding to NHS-labelled FeLV p27 under identical conditions. This lack of significant binding can be attributed to the weaker affinity demonstrated by #080 during earlier capture ELISA experiments, thereby reinforcing the reliability of our combined plate-based and solution-phase orthogonal methods for ranking mAb candidates. Additionally, the observed differences in apparent affinities and EC_50_ values for #080, as measured by detection ELISA versus in-solution SST, may be explained by either a non-native conformation of the FeLV p27 binding epitope when immobilized on the ELISA plate and/or by a specific interference caused by the NHS dye attached to primary amines in the specific epitope targeted by #080. Discerning which of the two factors explains said result was considered to be out of the scope of this work (see future works subsection below).

**Figure 4.**
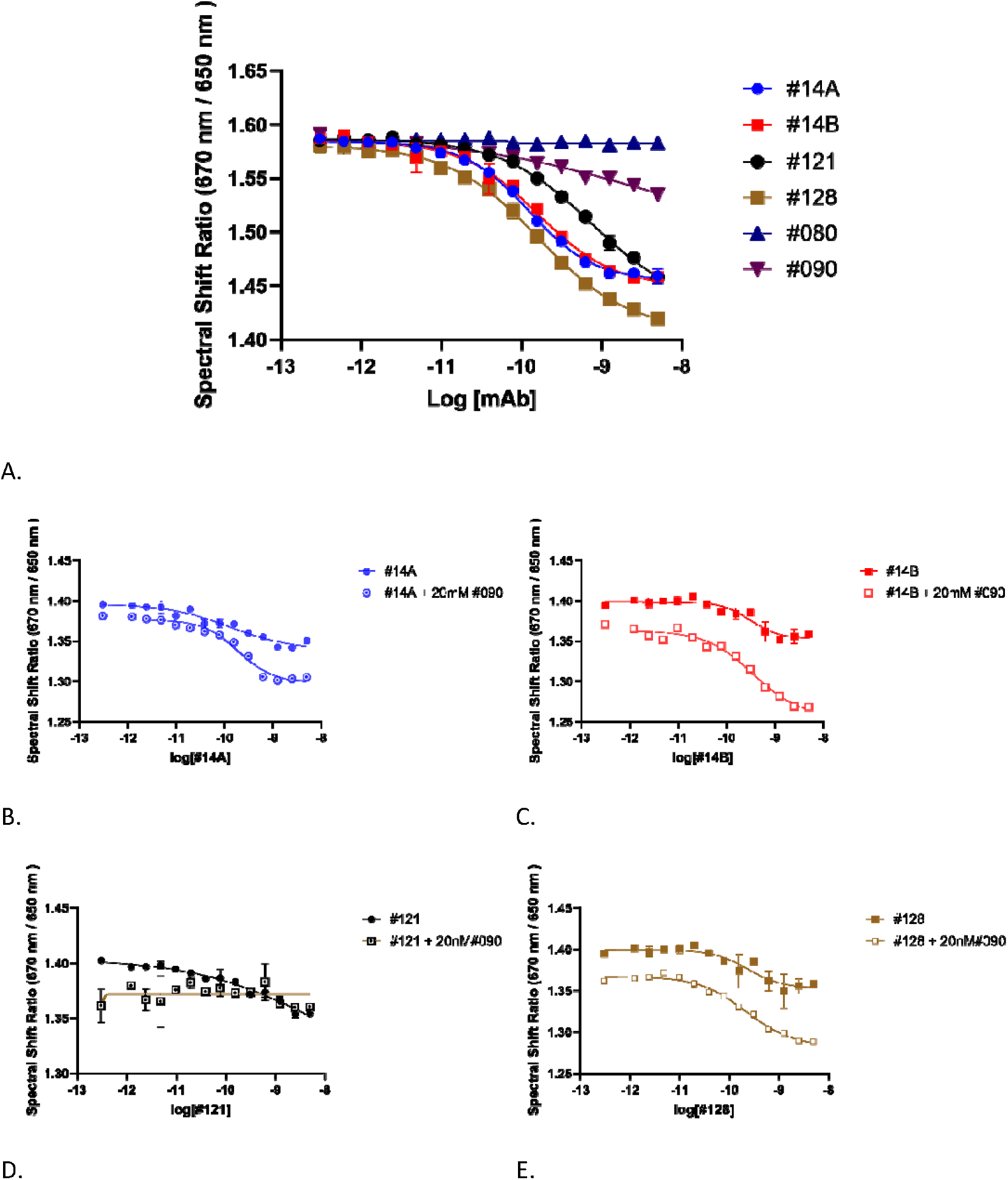
Spectral Shift Characterization of Monoclonal Antibody Affinities for FeLV p27. Spectral Shift assay characterization of FeLV mAbs binding to FeLV p27 antigen covalently labelled on primary amines with RED-NHS dye. A) Binary titrations, to further identify viable ELISA binders under standard conditions. Panels B) to E) Characterization of selected mAbs under stringent conditions to determine the most suitable for LFIA according to their ability to form ternary complexes with reference mAb #090.

Next, mAbs #14A-B, #121, and #128 were evaluated in SST antibody pairing assays to determine suitable pairs for sandwich assay development. This approach depends on effective ternary complex formation under stringent conditions, with one epitope of the NHS-labelled FeLV p27 antigen being saturated by reference mAb #090, which was selected to reduce uncertainties regarding the availability of another distinct, non-overlapping epitope (28). More specifically, a saturated FeLV p27–#090 complex (1:20 molar ratio) was titrated with each lead mAb in a dose-response format (Figures 4B-4E and Table 1). Said comparison enabled the classification of the mAb candidates in two groups according to their epitope binding properties: 1) candidates #14A-B, and #128, which presented fairly similar nanomolar K_d_s when binding to FeLV p27 alone or to the FeLV p27-#090 binary complex; and 2) #121, which managed to bind to FeLV p27 with a K_d_ of 4.64 nM, but did not show measurable binding to the FeLV p27-#090 complex in the nanomolar concentration range. Importantly, these results confirmed that candidates #14A-B, and #128 form an efficient ternary complex with FeLV p27-#090 binding to an immunogenic epitope that do not overlapps with #090’s epitope. In the same line, candidate #121 seems to compete with reference #090 for the same epitope (29,30). Hence, the antibody pairing SST methodology efficiently enabled us to narrow down the number of feasible LFIA sandwiches based on FeLV p27 to be tested for further selection of the novel mAbs to #14-#121, or #121-#128.

### 2.4. Immunochromatographic strip development for FeLV detection

2.4. Immunochromatographic strip development for FeLV detection.
| mAb<br>membrane<sup>a</sup> | mAb-AuNP conjugate<sup>b</sup> |  |  |  |  |  |  |  |  |  |  |  |  |  |  |
| --- | --- | --- | --- | --- | --- | --- | --- | --- | --- | --- | --- | --- | --- | --- | --- |
|  | #14A |  |  | #14B |  |  | #121 |  |  | #128 |  |  | #090 |  |  |
|  | D | N | P | D | N | P | D | N | P | D | N | P | D | N | P |
| #14A |  |  |  | - | - | - | - | - | w | - | - | - | - | - | m |
| #14B | - | - | - |  |  |  | - | - | w | - | - | - | - | - | m |
| #121 | - | - | m | - | - | m |  |  |  | - | - | s | - | - | - |
| #128 | - | - | - | - | - | - | - | - | w |  |  |  | - | - | w |
| #090 | - | - | m | - | - | m | - | - | - | - | - | s |  |  |  |

<sup>a</sup> Anti-FelV p27 mAbs were dispensed onto a nitrocellulose membrane at 0.3 mg/mL in PBS in a volume of 1 μL/cm. <sup>b</sup> mAb-AuNP conjugates were tested at an optical density (OD) of 4.0.

The LFIA platform architecture relies on a trinary interaction, where signal generation occurs through simultaneous binding of capture and detection antibodies to the target antigen. The orthogonal affinity-based antibody pairing method efficiently reduced potential LFIA combinations employing miniaturized assays that minimized material usage. However, a full evaluation of all four main mAbs and the best reference mAb in the LFIA format was conducted to determine how well results from the orthogonal affinity analysis aligned with a gold standard immunochromatographic strip method of higher translational value. For this, as described in Materials and Methods below, each mAb was dispensed on to a Hi-Flow plus 120 nitrocellulose membrane at a concentration of 0.3 mg/mL in PBS at a volume of 1 μL/cm in the Test line area (T), whereas each mAb-AuNP conjugate was dispensed on to the conjugate pad at an optical density (OD) of 4.0. Strip assembly was then carried out to ensure all 20 potential combinations, apart from self-paired configuration, and subsequently tested against antigen free sample diluent (D), a pool of negative FeLV serum samples (N) and a pool of low positive FeLV serum samples (P) (results shown in Table 2 and Figure S2).

In the absence of FeLV p27 antigen, all antibody pairing combinations showed no signal in the Test line when strips were evaluated against the sample diluent buffer or negative pool serum samples, confirming no non-specific binding, which could lead to false positives. On the contrary, according to the results shown above, significant signals were obtained for novel mAb pairs (membrane/conjugate): #14A/#121, #14B/#121, #121/#14A, #121/#14B, #121/#128 and #128/#121, which could be understood as a good correlation with the former SST analysis. LFIA experiments determined that mAb #121 showed similar activity, i.e.., stronger capture signals, to the reference mAb #090 when coated to the nitrocellulose membrane, whereas mAb #128 manifested the best detection performance in the series when conjugated to AuNP. Overall, the combination of novel mAbs #121/#128 was found to be the best performing pair in this format to validate our new methodology, compared with candidates #14A and #14B, which systematically led to weaker LFIA detection signals, because of which #14 mAbs were deprioritized.

At this point, it was deemed necessary to assess the universality of our novel methodology by comparing the results obtained with the classic gold-standard mAb pairing method, sandwich ELISA, at least with the most promising candidates. For this, peroxidase conjugation to best detection mAb #128 was carried out following the procedure described in the method section of this work and pairing compatibility with the novel antibody #121 and reference antibody #090 was assessed (Figure 5A). Similar to previous results obtained in mouse antiserum screenings (Figure S1), and LFIA screenings (Table 2, and Figure S2), antigen-free negative controls produced absorbance signals below a value of 0.1, meanwhile in the presence of 30 nM of the FeLV p27 antigen, Absorbance values increased proportionally with the amount of HRP-#128 conjugate tested, which suggested a specific detection signal, and matches with the trend observed by all the affinity assays performed along this work.

**Figure 5.**
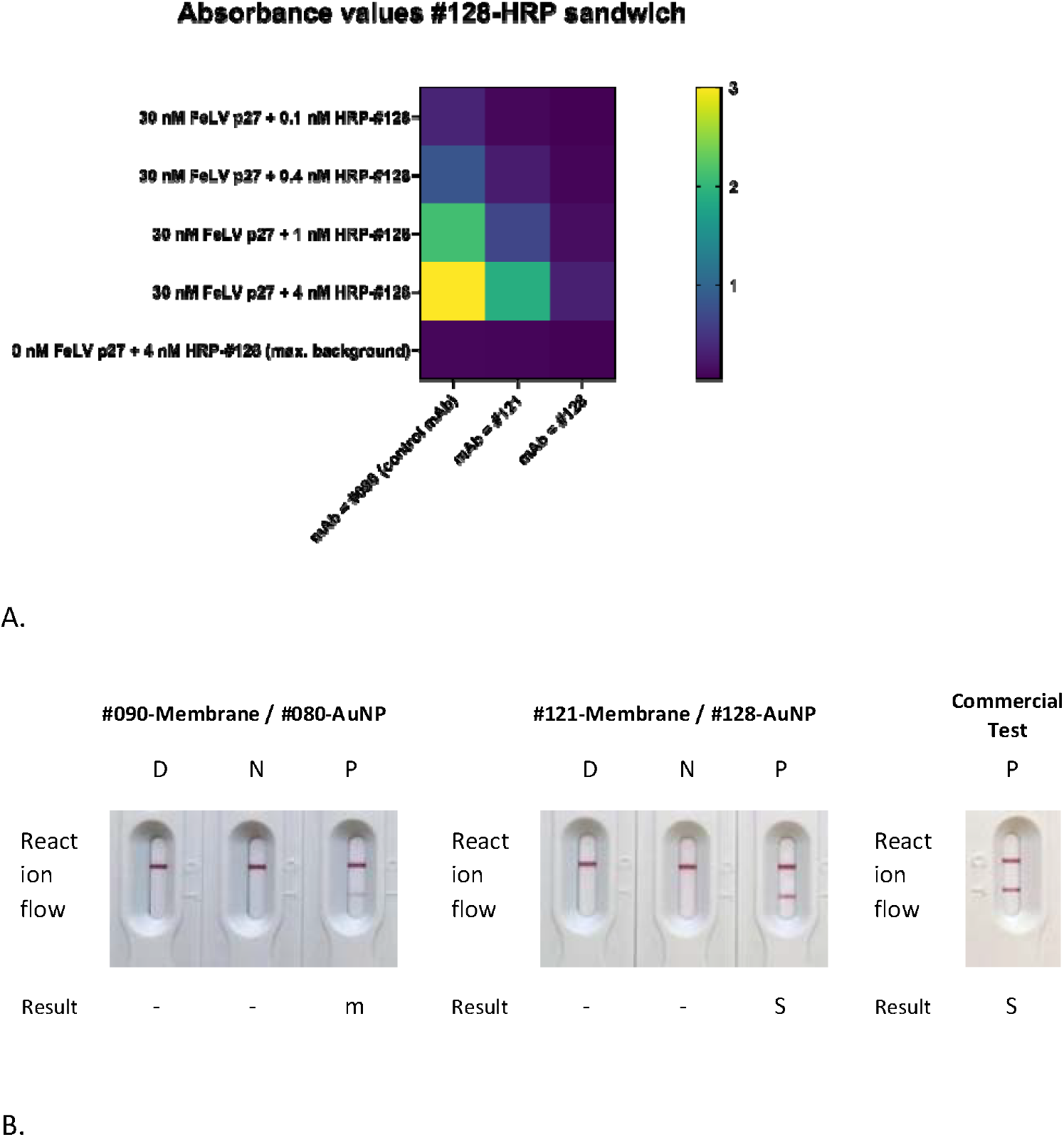
Translation of the binding studies towards a novel LFIA medical device based on #121 and #128 mAb candidates. A) Absorbance values at 450 nm obtained for each antibody in ELISA sandwich format against increasing concentrations of antibody #128-HRP conjugate in the absence of FeLV p27 antigen and in the presence of 30 nM of FeLV p27. B) LFIA screening of mAb pairs #121/#128 and #090/#080 at increased mAb membrane concentration (0.6 mg/mL) in comparison with a commercially available FeLV p27 detection strip. Results for each combination tested against sample diluent (D), negative FeLV serum pool (N) and low positive FeLV serum pool (P) are illustrated as negative (-), medium positive (m) and strong positive (S).

To further enhance the sensitivity of the LFIA system, and taking advantage of the sandwich ELISA results, a second LFIA test was carried out for the novel mAb pair #121/#128 and the reference pair #090/#080 employing an increased concentration of 0.6 mg/mL in a volume of 1 μL/cm for each membrane antibody. Consistently, the results shown in Figure 5B revealed no signal development when sample diluent and negative samples were assessed, while pair #121/128 showed an increased signal when dispensed at a higher concentration. Finally, the performance of both monoclonal antibody pairs was systematically evaluated against a commercially available LFIA for FeLV p27 detection. This comparison validated the efficacy of the newly developed test and demonstrated alignment with the innovative screening methodology adopted in this study, compatible with classic gold standard methods, like sandwich ELISA, that are more efficient when placed at the late stages of LFIA optimizations.

### 2.5. Conclusions and future works

The present study aimed to develop and evaluate new mAb pairings targeting a model antigen like FeLV p27 for LFIA applications, with the purpose of defining a new, low-cost, efficient methodology at finding alternatives to commercial antibody configurations. Using systematic affinity-based screening and mAb paring methods (14,29), including plate-based and in-solution SST assays of minimum sample consumption and straightforward implementation (7,31), the low nanomolar mAbs #121 and #128 were identified as compatible sandwiching mAbs targeting differential immunogenic epitopes of FeLV p27, and further tested in translational assays requiring more elaborated surface attachment protocols, or antibody labelling protocols, and higher amounts of materials. Indeed, #121 and #128 strong binding signals to FeLV p27 in both LFIA and sandwich ELISA formats were comparable to those of leading commercial pairs, with no observed non-specific binding, validating the whole innovative workflow proposed in the objectives of this work presented here. Furthermore, the applied methodology for FeLV p27 can be readily translated to other antigens of veterinary or clinical importance, particulary to current or emerging infectious diseases for which few reliable commercial LFIA tests are available, including Avian Influenza or African Swine Fever Virus, as well as human pathogens like *Helicobacter pylori* or Mpox Virus, among others. By reducing traditional antibody pairing screening procedures in early stage development, this approach could increase the availability of diagnostic tools essential for disease control and prevention.

In the scope of our proof of concept with FeLV p27, further adjustments, such as increased membrane mAb concentration, enhanced the sensitivity, of the #121/#128 pairing in the LFIA prototype. As such, the main methodologic objectives of our research, e.g. identification of high-performing novel mAbs pairs, elimination of non-specific responses, and demonstration of effective LFIA translation, were successfully met. The results are a proof of concept that the use of the new mAbs in LFIA are suitable for FeLV p27 detection and should be applicable in diagnostic contexts.

Future research will include detailed structural studies and epitope identification (32), utilizing FeLV p27 mutants especially focusing on its central and C-terminal domains (16,18), which could be a great starting point to refine antibody #121 and address the limitations observed in tests like SST in the stringent format, in which the binding signal was weaker and had led to less defined top and bottom plateaus, suggesting further optimization could be done to enhance the sensitivity of the final LFIA test.

Additional kinetic analyses, such as measurement of association and dissociation rates (K_on_ and K_off_ (33), which were not included in this study, could also be incorporated in subsequent SPR studies. Although prototype results suggest that the binding kinetics of selected candidates are compatible with LFIA, a more comprehensive understanding of these parameters by SPR will help optimize antibody #121 and advance antibody #128 and its conjugates. Incorporating these research directions with current findings is expected to support the ongoing development of diagnostic solutions for FeLV p27 detection.

## 3. Materials and Methods

### 3.1. Materials and equipment

FeLV p27 protein (cat. No. RAG0078) and monobiotinylated FeLV p27 protein (cat. No.RAG0078BIOT) was acquired from Rekom Biotech (Granada, Spain). Reference antibodies #090 (cat. No. 1903090) and #080 (cat. No. 1803080) were purchased from Biosynth Ltd (United Kingdom, https://www.biosynth.com/p/CZ9111/feline-leukemia-virus-mouse-monoclonal-antibody). Anti-Mouse IgG (H+L)-Peroxidase antibody produced in Rabbit (cat. No. SAB3701083), Streptavidin from *Streptomyces avidinii* (cat. No. S4762) and Horseradish peroxidase (HRP, cat. No. 31490, activity: 279 U/mg) was obtained from Sigma/Aldrich (Madrid, Spain). Molecular Probes™ Streptavidin, horseradish peroxidase conjugate (cat. No. 10267002) was purchased from ThermoFisher Scientific (Madrid, Spain).

SP2/O-Ag14 non-secreting Ig Murine Cell Line was provided by UGR (Universidad de Granada, Granada, Spain). Female BALB/c mice (BALB/cAnNCrl) were supplied by Charles Rivers Laboratories (L’Arbresle, France) and cared for by the Animal Experimentation Unit of the López Neyra Institute of Parasitology and Biomedicine (IPBLN–CSIC; Granada, Spain) in compliance with the European Directive 2010/63/EU and the Spanish laws and guidelines (RD53/2013) concerning the protection of animals used for experimentation and other scientific purposes.

Freund’s adjuvant (Complete, cat. No. F5881; Incomplete, cat. No. F5506), Dulbecco’s Modified Eagle’s Medium – high glucose (DMEM, cat. No. D6546), Gentamicin solution 50 mg/mL (cat. No. G1397), MEM Non-essential Amino Acid solution 100x (cat. No. M7145), Polyethylene glycol solution 50% (w/v) Hybri-Max (PEG 1500, cat. No. P7181), Fetal Bovine Serum (cat. No. F7524), Hybridoma Fusion and Cloning Supplement 50x (HFCS, cat. No. 11363735001), Red blood cell lysing buffer Hybri-Max (cat. No. R7757) and L-Alanyl-L-Glutamine 200 mM (cat. No. G8541) was obtained from Sigma/Aldrich (Madrid, Sapin). HT supplement (hypoxanthine 5 mM, thymidine 0.8 mM, cat. No. 41065-012) and HAT supplement (HT supplemented with aminopterin 20 μM, cat. No. 21060-017) were purchased from ThermoFisher Scientific (Madrid, Spain). Dimethyl sulfoxide (DMSO) Cell culture grade (cat. No. A3672) was provided by PanReac AppliChem (Barcelona, Spain).

TMB One Component HRP Microwell Substrate was provided by Surmodics IVD, Inc (Eden Prairie, Minnesota, United States). Sodium periodate ≥98% (cat. No. 013798.09) and Sodium borohydride, 98% (cat. No. 013432.22) was purchased from ThermoFisher Scientific (Madrid,Spain).

Flat-bottom 96-well polystyrene ELISA plates (cat. No. 655101) and EASYstrainer™ 70 μm Sterile meshes (cat. No. 15391067) for culture medium filtration were purchased from Greiner BIO-ONE (Madrid, Spain). HiPrep™ Desalting columns with Sephadex G-25 resin (cat. No. 17508701), HiTrap® Protein G High Performance (cat. No. 17040501) and Vivaspin® 500, 3 kDa MWCO Polyethersulfone (cat. No. 28932218) were obtained from Cytiva Life Sciences Solutions (Wilmington, United States).

ELISA plate absorbance was registered by a BioTek 800 TS absorbance reader from Agilent Technologies (Madrid, Spain). Microplate washing was carried out using a RT-3900 automatic 96-well microplate washer from Biogen Científica. Cell cultures were maintained at 37 ^°^C and 5% CO_2_ in a HF-212UV Heal Force incubator also obtained from Biogen Científica. Antibody purification was preformed employing a NGC Quest 10 Plus Chromatography System #7880003 from Bio-Rad.

Spectral shift analysis was carried out employing a Dianthus-uHTS platform by NanoTemper Technologies®.

### 3.2. Antibody production and ELISA characterization

#### 3.2.1. Monoclonal antibody production

For the generation of specific mAbs against the FeLV p27 protein, two-month-old female Balb/c mice were administered three doses of 100 μg of the recombinant FeLV p27 antigen in a 200 μL 1:1 (v/v) emulsion of 0.22 μm filtered phosphate buffer and either complete Freund’s adjuvant for the first dose or incomplete for the subsequent boosts. Upon completion, a final inoculation in the absence of Fruend’s adjuvant was carried out 21 days after the final boost and four days prior to hybridoma production.

Hybridoma production was carried by isolating the B cells from the spleen of immunized mice post-sacrifice and subsequently fusing these cells with SP2/0-Ag14 myeloma cells at a four to one ratio. Cells were then plated at a density of 150.000 cells per well and incubated for 24 hours before selection medium containing HAT supplement was added. Cell cultures were maintained at 37 ^°^C / 5% CO_2_ for 10 to 12 days.

Screening of the hybridoma culture supernatant for anti-FeLV P27 antibodies was carried out by detection ELISA. For this, 50 μL of each supernatant was diluted with one volume of PBST and then added to plates coated with 0.5 nM of the FeLV p27 protein. Wells exhibiting an absorbance signal value greater than 1.0, were then reevaluated by checkerboard ELISA and selected hybridomas were then cloned. Upon confirming hybridoma monoclonality, cell culture expansion was carried out to obtain sufficient mAb for assay development.

Monoclonal antibody isolation from culture medium was carried out by eluting each supernatant through a Cytiva HiPrep desalting column with Sephadex G-25 resin with phosphate buffer, followed by a 0.45 μm microfiltration. Antibodies were then purified using a Cytiva HiTrap protein G HP column and gathered fractions were once again passed through a HiPrep desalting column. Purified antibodies were then quantified and stored at -20 ^°^C in 20 mM pH 7.0 phosphate buffer containing 0.15 M of NaCl, 5% Trehalose and 0.1% sodium azide.

#### 3.2.2. Anti-P27 FeLV monoclonal antibody characterization by ELISA

##### Capture ELISA

Microtiter plates were coated with 100 μL of each anti-FeLV p27 mAb at a concentration of 7.14 nM in PBS. Following an overnight incubation at 4 ^°^C, the plates were then washed four times with 200 μL of PBST. Antibody capture was carried out by adding 100 μL of a sixteen-point two-fold serial dilution of the biotinylated FeLV p27 antigen at a starting concentration of 12 nM. After a one-hour incubation, the plates were then once again washed and 100 μL of peroxidase-labelled streptavidin was added to each well and incubated at room temperature for another hour. After a final plate washing step, 100 μL of enzyme substrate was added to each well and incubated for 15 minutes at room temperature. Upon completion, 100 μL of stop solution was added and the absorbance was promptly registered, which was fitted to a four-parameter logistic model in GraphPad Prism 10 for EC_50_ determination. The EC_50_ value was determined by fitting the obtained absorbance values to a four-parameter logistic (4PL) model, y = d + (a -d) / (1 + (x/c)^b), using GraphPad Prism 10. In this equation, a represents the top asymptote (maximal signal), d the bottom asymptote (minimal signal), b the Hill slope (which indicates the steepness of the curve), and c the inflection point, representing the concentration at which half the maximal signal is reached. The EC_50_ value corresponds to parameter c, reflecting the ligand concentration required to occupy 50% of the available binding sites and thereby characterising the apparent affinity of the antibody-antigen interaction under the assay conditions (23).

##### Detection ELISA

Microtiter plates were coated overnight at room temperature with 100 μL of streptavidin in PBS at a concentration of 19.2 nM. After plate washing, 100 μL per well of a sixteen-point two-fold serial dilution of the FeLV p27 antigen at a starting concentration of 12 nM was added and incubated for one hour at room temperature. Antigen detection was performed by incubating the different anti-FeLV p27 mAbs to the previously washed plates for one hour at room temperature. After a final plate washing step, 100 μL of the enzyme labelled secondary antibody was added at a 10000-fold dilution in PBST and incubated for one hour at room temperature. Signal development was carried out by adding 100 μL of enzyme substrate and incubating for 15 minutes before adding 100 μL of stop solution and recording the obtained absorbance, which were fitted to the same four-parameter logistic model in GraphPad Prism 10 for EC_50_ determination and comparison with corresponding EC_50_ values obtained by Capture ELISA.

##### Sandwich ELISA

Wells were coated with 100 μL of each mAb at a concentration of 7.14 nM in PBS. After an overnight incubation at 4 ^°^C, plates were subsequently washed and 100 μL of the FeLV p27 antigen was added at a concentration of 24 nM in PBST for one hour at room temperature. Plates were then once again washed before adding 100 μL per well of the antibody #128 □ peroxidase conjugate and incubating for a further hour at room temperature. Lastly, upon washing, 100 μL of the enzyme substrate was added for 15 minutes, following the addition of 100 μL of stop solution and plate reading at 450 nm to monitor correct sandwich formation.

#### 3.2.3. Monoclonal antibody isotyping

The Isotope of all generated antibodies was determined using a Rapid ELISA mouse mAb Isotyping Kit by Invitrogen (cat. No. 37503; ThermoFisher Scientific) following the manufacturer’s instructions.

#### 3.2.4. Monoclonal antibody peroxidation

Following the periodate oxidation method, antibody conjugation to HRP was carried out by adding sodium periodate (101.4 μL, 0.1 M, 0.6 equivalents) to 0.5 mL of HRP (1000 U, 279 U/mg) in water. The latter mixture was stirred in darkness for 20 minutes at room temperature and remaining reactants were separated using Vivaspin 500 3 kDa MWCO filters. Five cycles of 15 minutes at 13300 rpm were carried out using 1 mM sodium acetate buffer pH 4.4, yielding a final volume of 0.5 mL. Upon completion, pH neutralization was carried out by adding 8.3 μL of 0.1 M carbonate buffer pH 9.5. Subsequently, 62.5 μL of the activated HRP was added to 0.1 mL of the mAb #128 at 2.5 mg/mL in 0.1 M carbonate buffer pH 9.6 and stirred for two hours at room temperature. After incubation, 1/20 volume of sodium borohydride at 5 mg/mL was added and the mixture was stirred for 30 minutes at 4 ^°^C, before a further 1/10 volume of sodium borohydride was added and stirred for another 30 minutes. Upon completion, reactant elimination and buffer exchange were carried out using Vivaspin 500 3 kDa MWCO filters (five 15-minute cycles at 13300 rpm). The obtained conjugate was used without further purification.

### 3.3. Spectral shift analysis

#### 3.3.1. Protein labelling

Primary amines of the FeLV p27 antigen were covalently labelled with Protein Labelling Kit RED-NHS 2nd Generation (cat. No. M0-L011; NanoTemper Technologies®) by adding 300 µM NHS dye to 90 µL of 10 µM FeLV p27 protein, then incubating for 30 minutes at room temperature in the dark. Unreacted dye was removed via the kit’s B-Column. Protein concentration and labelling efficiency (>70%) were confirmed by UV-vis spectroscopy according to manufacturer’s recommendations.

#### 3.3.2. Binary interaction analysis

Labelled antigen and unlabelled mAbs were prepared at 2x concentrations in binding buffer (PBS, pH 7.4, 0.1% DDM), mixed 1:1, and incubated for 60 minutes at room temperature. For binary interactions, 5 nM labelled antigen and a 15-point two-fold serial dilution of mAb (starting at 50 nM) were used as final concentrations, and data obtained employing a Dianthus-uHTS platform by NanoTemper Technologies® were fitted to the four-parameter logistic model in GraphPad Prism 10 for K_d_ determination. In this case, the equation of the 4PL model (y = d + (a - d) / (1 + (x/c)^b), a represents the top asymptote (maximal signal), d the bottom asymptote (minimal signal), b the Hill slope (which indicates the steepness of the curve), and c the inflection point, representing the concentration at which half the maximal signal is reached. The K_d_ value corresponds to parameter c, reflecting the ligand concentration required to occupy 50% of the available binding sites and thereby characterising the apparent affinity of the antibody-antigen interaction under the assay conditions, and for which a simple 1:1 stoichiometry is assumed for a more direct comparison with ELISA’s EC_50_ values reporting for apparent affinity (23).

#### 3.3.3. Stringent assay

For stringent assays, 1 nM labelled antigen previously saturated with 20 nM reference antibody #090 identified top mAb candidates for ternary complex formation. All experiments were repeated; fluorescence emission ratios (670 nm/650 nm) at 25 ^°^C indicated FeLV p27 binding, employing a Dianthus-uHTS platform by NanoTemper Technologies® were fitted to a four-parameter logistic model in GraphPad Prism 10 for K_d_ determination as described above.

### 3.4. Lateral flow immunoassay development

#### 3.4.1. Colloidal gold nanoparticle (AuNP) labelling with anti-FeLV p27 monoclonal antibodies

##### Conjugation buffer pH optimization

First, the optimal pH for AuNP conjugation of each anti-FeLV p27 mAb was determined by adding a serial dilution of K_2_CO_3_ 0.1 M to reactions tubes containing 1 mL of 40 nm AuNP (purchased from BBI Solutions UK, cat. No. EM.GC40 SPL). Then, 10 μg of mAb were added to each reaction tube and, after a 10-minute incubation, 100 μL of NaCl 10% were added and the mixture was stirred for an additional 5 minutes. Each solution was then measured at 520 nm and final conjugation buffer pH values were selected based on conjugate stability, falling within a range of 8.0 – 8.8 for all six mAbs.

##### Conjugation procedure

Conjugation was carried out by adjusting the pH of 10 mL of 40 nm AuNP with K_2_CO_3_ 0.1 M followed by the dropwise addition of 100 μg of each mAb. The latter solution was then stirred for 30 minutes before adding one volume of 0.5% BSA blocking solution. Upon stirring for an additional 30 minutes, each mixture was centrifuged (6000 g, 30 minutes) and the resulting pellet was resuspended in 1 mL of borate buffer 10 mM, BSA 0.5% and sucrose 15%. Conjugates were then stored at 4 ^°^C.

#### 3.4.2. Anti-FeLV p27 monoclonal antibody conjugate dispensing

Each mAb-AuNP conjugate was diluted with Borate buffer 10 mM, BSA 0.5% and Sucrose 15% to an OD of 4.0 and then spayed onto 8 mm next generation hydrophilic polyester conjugate pads (purchased from Ahlstrom, cat. No. 6614) until full saturation. Conjugate pads were dried at 37 ^°^C for 1 h and then stored in aluminium foils with desiccants at room temperature.

#### 3.4.3. Monoclonal antibody binding to nitrocellulose membrane

For membrane dispensing, each mAb was diluted to a concentration of 0.3 mg/mL or 0.6 mg/mL in PBS and sprayed at a speed of 1 μL/cm onto a Hi-Flow Plus 120 nitrocellulose membrane (purchased from Sigma-Merck, cat. No. SHF1200425) in the position of the Test line (T). In the position of Control line (C), goat anti-mouse IgG 0.5 mg/mL was also sprayed in a volume of 1 μL /cm. Lastly, membranes were dried at 37 ^°^C for 2 h and stored in aluminium foils with desiccants at room temperature.

#### 3.4.4 Immunochromatographic strip assembly

Briefly, strip assembly for each mAb was constructed by placing onto a PVC backing card a cotton-based blood separator sample pad, conjugate pad containing the previously dispensed mAb-AuNP conjugate, nitrocellulose membrane with dispensed Test and Control lines and a cotton based hygroscopic absorbent pad. Assembled cards were then cut into 4 mm strips and housed in independent cassettes.

#### 3.4.5 FeLV analysis by LFIA

Screening assays were performed out by adding 10 μL of FeLV negative or positive feline serum onto the sample pad, immediately followed by the addition of 2 drops of borate buffer-based sample diluent. Sample migration was carried out for 10 minutes and, upon completion, visual interpretation of results was conducted, considering two red lines as positive, one control line and no test line as negative and no control line as invalid.

## Acknowledgements

Authors gratefully appreciate the excellent technical support provided in antibody purification by Nidia Martí Jiménez and María José Barroso Montoro.

## Author contributions

**Hadyn Duncan:** Methodology, Conceptualization, Investigation, Visualization, Writing – original draft, English, Project Management. **Elisabeth Domingo-Contreras:** Methodology, Formal analysis. **Sotirios Athanasiou:** Methodology. **Rosario Fernández-Godino:** Writing – review & editing, Visualization, Funding Acquisition. **Francisco Castillo:** Methodology, Conceptualization, Investigation, Visualization, Writing – original draft, final manuscript – review & editing, Project Management, Supervision, Funding Acquisition. **Ana Camacho:**Methodology, Conceptualization, Investigation, Visualization, Writing –final manuscript–, review & editing, Supervision, Funding Acquisition.

## Additional information

Monoclonal antibodies #121 (cat. No. MAB0001) and #128 (cat. No. MAB0002) for FeLV p27 reported herein are available for purchase at Rekom Biotech (Granada, Spain).

## Supplementary information

**S1.**
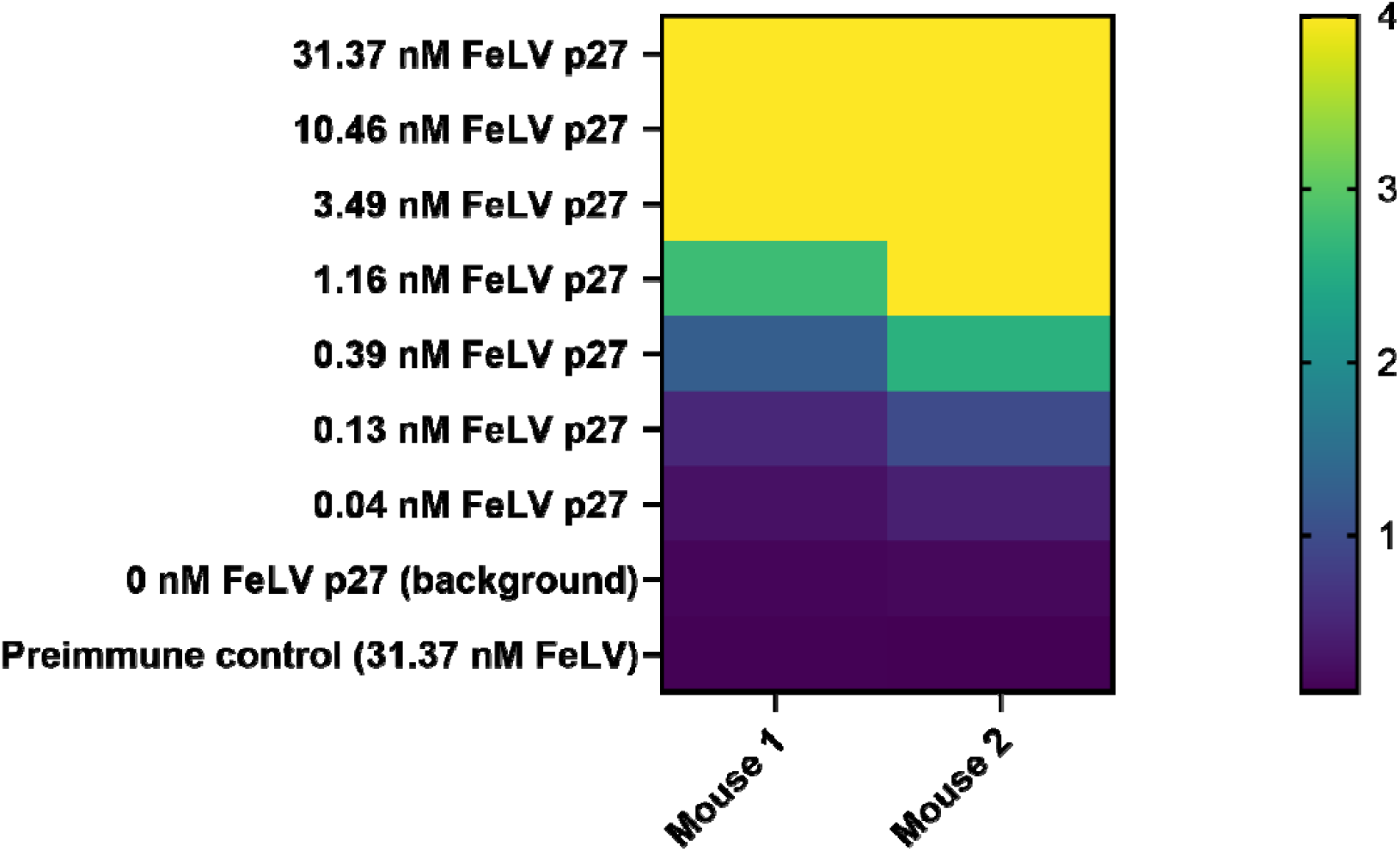
Absorbance values obtained from mouse antiserum screening in detection ELISA. Microtiter plates were coated overnight at room temperature with 100 μL of streptavidin in PBS at a concentration of 1 μg/mL. After plate washing, 100 μL per well of an eight-point three-fold serial dilution of the biotinylated FeLV p27 antigen at a starting concentration of 31 nM was added and incubated for one hour at room temperature. Antigen detection was performed by incubating the obtained anti-FeLV p27 serum at 1000-fold dilution to the previously washed plates for one hour at room temperature. After a final plate washing step, 100 μL of the enzyme labelled secondary antibody was added at a 10000-fold dilution in PBST and incubated for one hour at room temperature. Signal development was carried out by adding 100 μL of enzyme substrate and incubating for 15 minutes before adding 100 μL of stop solution and recording the obtained absorbance.

**Figure S2.**
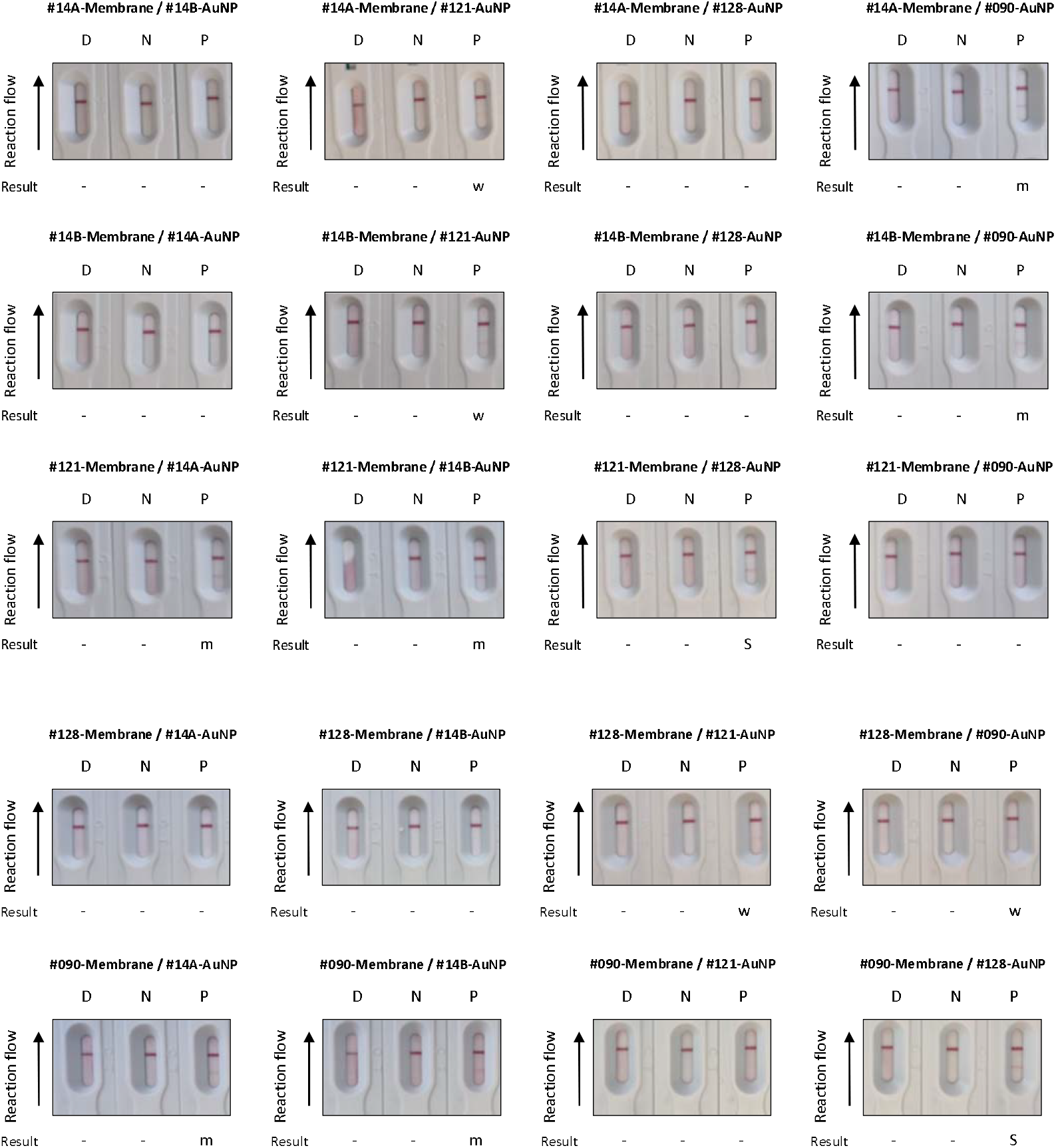
Visual representation of LFIA strips results obtained during antibody pairing screening. Results for each combination tested against sample diluent (D), negative FeLV serum pool (N) and low positive FeLV serum pool (P) are illustrated as negative (-), weak positive (w), medium positive (m) and strong positive (S).

