## Supplementary Figures 1 and 2 for "Affinity-based selection of anti-Feline Leukaemia Virus p27 monoclonal antibodies for efficient lateral flow assay development"

**Supplementary information**

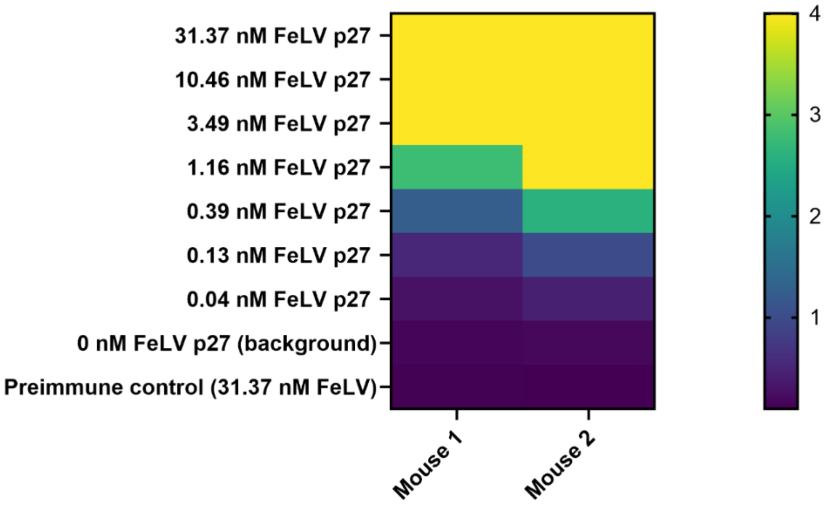


***S1. Absorbance values obtained from mouse antiserum screening in detection ELISA.*** Microtiter plates were coated overnight at room temperature with 100 µL of streptavidin in PBS at a concentration of 1 µg/mL. After plate washing, 100 µL per well of an eight-point three-fold serial dilution of the biotinylated FeLV p27 antigen at a starting concentration of 31 nM was added and incubated for one hour at room temperature. Antigen detection was performed by incubating the obtained anti-FeLV p27 serum at 1000-fold dilution to the previously washed plates for one hour at room temperature. After a final plate washing step, 100 µL of the enzyme labelled secondary antibody was added at a 10000-fold dilution in PBST and incubated for one hour at room temperature. Signal development was carried out by adding 100 µL of enzyme substrate and incubating for 15 minutes before adding 100 µL of stop solution and recording the obtained absorbance.

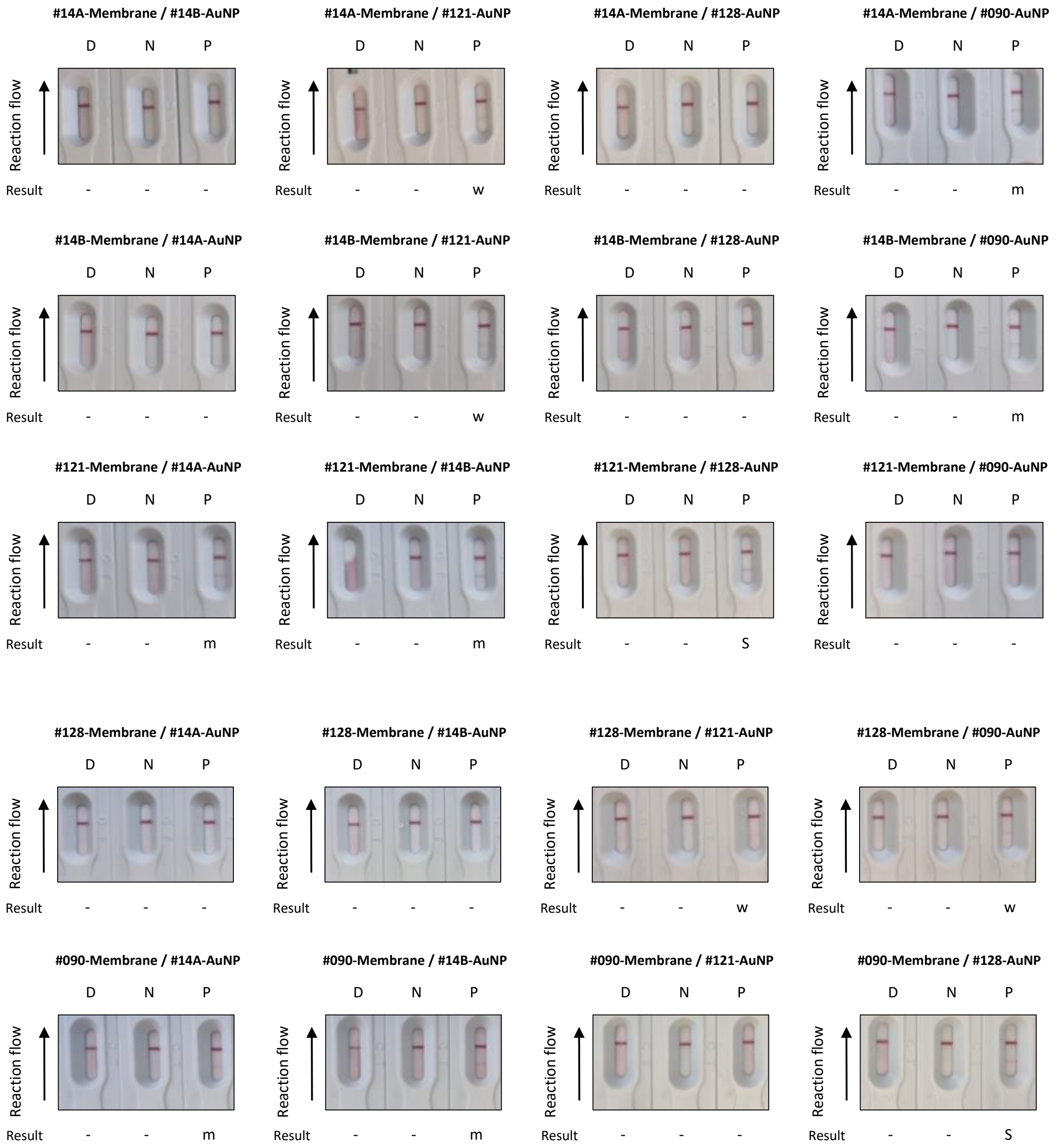


***Figure S2.*** **Visual representation of LFIA strips results obtained during antibody pairing screening**. Results for each combination tested against sample diluent (D), negative FeLV serum pool (N) and low positive FeLV serum pool (P) are illustrated as negative (-), weak positive (w), medium positive (m) and strong positive (S)
